# Angiotensin converting enzyme 2 (ACE2) is expressed in murine cutaneous under single-cell transcriptome resolution

**DOI:** 10.1101/2021.10.07.463533

**Authors:** Chenyu Chu, Chen Hu, Li Liu, Yuanjing Wang, Yili Qu, Yi Man

**Author notes:** **Corresponding author**: **Yili Qu, DDS, PhD**, Associate Professor, State Key Laboratory of Oral Diseases & National Clinical Research Center for Oral Diseases, West China Hospital of Stomatology, Sichuan University, 14#, 3rd, section, Renmin South Road, Chengdu 610041, China, **Yi Man, DDS, PhD**, Chair and Professor, State Key Laboratory of Oral Diseases & National Clinical Research Center for Oral Diseases & Department of Oral Implantology, West China Hospital of Stomatology, Sichuan University. These authors contributed equally to this work (Chenyu Chu, Chen Hu).

## Abstract

Angiotensin converting enzyme 2 (Ace2) is widely distributed in human organs, which was identified as a functional receptor for severe acute respiratory syndrome (SARS) coronavirus in human beings. It was also confirmed that SARS-CoV-2 uses the same cell entry receptor, ACE2, as SARS-CoV. However, related research still not discover the expression data associated with murine skin under single cell RNA resolution. In this study, we performed single-cell RNA sequencing (scRNA-seq) on unsorted cells from mouse dorsal skin after 7 days post-wounding. 8312 sequenced cells from four skin samples met quality control metrics and were analyzed.

## 1. Introduction

Angiotensin converting enzyme 2 (Ace2) is widely distributed in human organs, especially cardio-renal tissues, the digestive systems and respiratory tract. [1-4] It functions as a membrane-associated extracellular protein that cleaves angiotensin (Ang) I to generate Ang 1-9.[2] Later, Ace2 was identified as a functional receptor for severe acute respiratory syndrome (SARS) coronavirus [5, 6]. In mouse, SARS-Cov Spike protein bind to Ace2, resulting in aggravated acute lung failure through deregulation of the renin-angiotensin system. In the recent 2019 coronavirus epidemic (COVID-19) caused by a novel virus SARS-CoV-2, 2129 people had died by 20 February. Infections worldwide have topped 75000; more than 74000 are in China.[7] It was confirmed that SARS-CoV-2 uses the same cell entry receptor, ACE2, as SARS-CoV.[8] To explore the expression of Ace2 in mouse skin wound model and potential functional roles of it, we performed single-cell RNA sequencing (scRNA-seq) on unsorted cells from mouse dorsal skin after 7 days post-wounding. 8312 sequenced cells from four skin samples met quality control metrics and were analyzed.

## 2. Method and materials

### 2.1 Experimental model

All animal procedures were approved by the Institutional Ethics Committee of Sichuan University. Animals included C57BL/6 male mice at ages from 6 to 8 weeks. Hair at the surgical area was removed by plucking against the growth direction. Full-thickness circular excisional wound (diameter=6mm) was created at the dorsal skin of mice. A sterile Tegaderm film (3M) was placed above the wound to protect the wound area. Then annular silicone splints (inner diameter=8mm, outer diameter=12mm) were sutured with the Tegaderm film and underlying skin in order to minimize the contraction of the dorsal muscle. After healing for a week, mice were euthanized for sample harvest.

### 2.2 Specimen harvest

We obtained skin samples by cutting off skin at the wound area (circular, diameter=10mm). Subcutaneous tissues were removed. A total of four tissues were harvested. The tissues were washed in a 100 mm petri dish containing 20 ml of phosphate-buffered saline (PBS). Then they were transferred to a 50 mm petri dish containing 100μL of Enzyme G (Epidermis Dissociation Kit mouse, Miltenyi) and 3.9 ml of PBS buffer with the dermal side facing downwards. Tissues were digested for 16 hours at 4°C. Then they were transferred into a 50 mm petri dish containing 4mL of 1×Buffer S (Miltenyi). Epidermis was peeled off from the skin using tweezers, and was cut into pieces. Enzyme mix containing 3.9 ml of 1×Buffer S, 100μl of Enzyme P, and 20μl of Enzyme A (Miltenyi) stored in a gentleMACS™ C Tube was used to digest the epidermis pieces for 20 minutes at 37□. Stop enzymatic reaction by adding 4 ml of PBS that contain 0.5% bovine serum albumin (BSA). A gentleMACS dissociator (Miltenyi) was applied to automatically dissociate the epidermis (Program B). The sample was passed through a 70μm cell strainer (Corning), centrifuged at 300×g for 10 minutes at room temperature, and suspended with PBS that contain 0.5% BSA. Cells were gently washed twice and stored in an ice box. For the dermis part, they were first cut into pieces (diameter < 1mm). The tissue was mixed with 10ml enzyme mix containing Type I Collagenase (3125U/Ml, Gibco) and 2.5ml trypsin (Gibco), and poured into a gentleMACS™ C Tube. After dissociating the tissue on gentleMACS dissociator for 37s (Skin mode), another 10ml enzyme mix was added. The sample was digested for 2.5 hours at 37□ in a rotary machine (Peqlab). Then the dermis sample was passed through a 70μm cell strainer (Corning), centrifuged at 300×g for 5 minutes at room temperature, and suspended with 3ml red blood cell lysis buffer (Solarbio). After 3 minutes, the cell suspension was centrifuged and gently suspended with RPMI 1640 medium (Hyclone). Cells were gently washed twice with PBS containing 0.5% BSA and stored in an ice box. The epidermis and dermis cell solutions were mixed together as a whole. The sample was centrifuged, and suspended with 100μl Dead Cell Removal MicroBeads (Miltenyi). After incubation for 15min at room temperature, the cell suspension was diluted in 3ml 1× Binding buffer (Miltenyi). LS columns (Miltenyi) were used for removal of dead cells and debris. The negatively selected live cells pass through the column, and were suspended with PBS containing 0.05% BSA. Finally, we proceeded with the 10x Genomics® Single Cell Protocol.

### 2.3 Single-cell encapsulation and library generation

Single cells were encapsulated in water-in-oil emulsion along with gel beads coated with unique molecular barcodes using the 10× Genomics Chromium Single-Cell Platform. For single-cell RNA library generation, the manufacturers’ protocol was performed. (10×Single Cell 3’ v3) Sequencing was performed using a NovaSeq PE150 mode with 94574 reads per cell. The Cell Ranger software was used to align reads and generate expression matrices for downstream analysis.

## 3. Results

Unsupervised clustering using Seurat categorized the cells into 21 clusters based on global gene expression patterns. Using the differentially expressed gene signatures, we annotated these clusters (Fig. 1a). 10% cells from wounded skin express Ace2 (Fig.1b), the majority of which belong to keratinocytes (cluster 2, marker genes: Krt1, Krt10, Loricrin) in the epidermis (9) and sebocytes in fig. 2 (cluster 12, marker genes: Mgst1, Scd1, Dhcr24, Elovl6) (10). We also analyzed bulk RNA-Seq profiles of mouse healthy dorsal skin. ACE2 was expressed in healthy skin as well (Fragments per kilobase million (FPKM) values more than 1). According to Gene Ontology (GO) analysis, cluster 2 (keratinocyte) was enriched for genes related to keratinocyte differentiation, skin development, epidermis development, establishment of skin barrier, and keratinization (Fig. 3). The epidermis is a stratified epithelium composed of several layers of keratinocytes, which provides a physical barrier between the environment and the organism, thereby protecting it from external agents and pathogens, and limiting the loss of fluids.[9] Cornification is achieved by keratinocytes passing through 4 cell layers of the epidermis: the stratum basale, the stratum spinosum, the stratum granulosum (SG), and the stratum corneum in fig.4 (SC) [10]. Interestingly, Ace2+ keratinocytes (44%) in cluster 2 is featured by up-regulated expression of keratins, e.g. Krt10, Krt78, which are markers for differentiated keratinocytes located at suprabasal layers of epidermis [9]. While Ace2-keratinocytes (56%) are featured by high expression level of Late cornified envelope group I (LCE1) genes, filaggrin2 (Flg2) and Loricrin (Fig. 5), the encoded protein of which are key elements for cornified cell envelope (CE). As SG keratinocytes transition into SC, the keratinocytes become flattened and denucleated (which are then called corneocytes). Lamellar bodies (containing extracellular components, e.g. lipids, corneodesmosin, and kallikreins) are secreted into the intercellular space between the corneocytes and fill with lipid. These structures are often described as bricks (corneocytes) and mortar (intercellular lipids), which together provide a highly hydrophobic barrier, namely cornified cell envelope (CE), against the environment [10-12]. The gene expression profiles of Ace2+ and Ace2-cells indicate that keratinocytes in the spinous and granular layers of epidermis express Ace2, while corneocytes at the outermost cornified layer do not. GO analysis of cluster 12 (sebocytes) exhibited enriched gene expression related to cellular respiration, ATP metabolic process and oxidative phosphorylation (Fig. 3). Cellular respiration, through oxidative phosphorylation (OXPHOS), constitutes the main oxygen-consuming and ATP-producing processes [13].

**Fig. 1.**
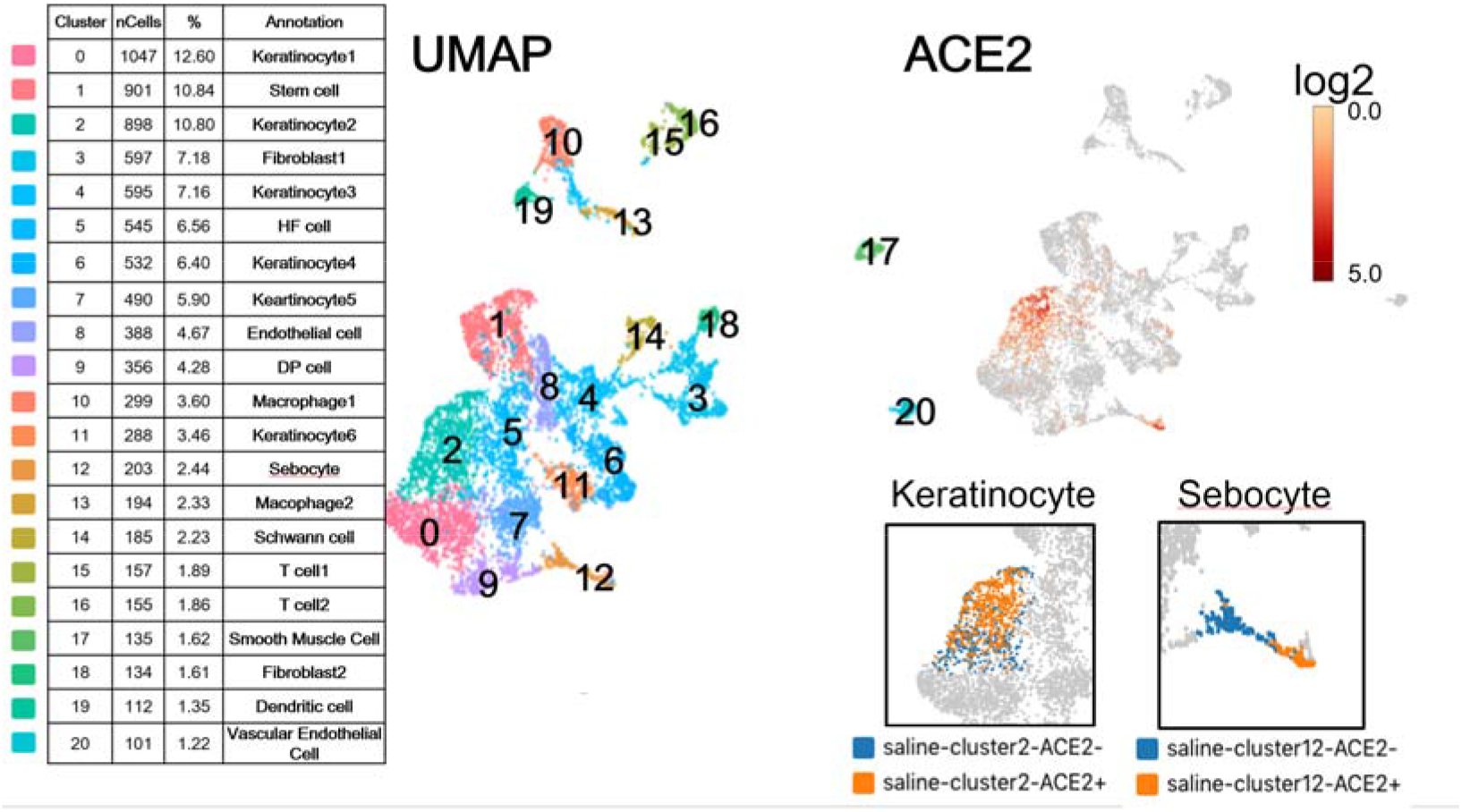
Clustering ACE2-positive area in UMAP, further marked related cells as keratinocyte and sebocyte, which express highly ACE2 level in murine full-thickness skin wound healing process at day 7.

**Figure 2.**
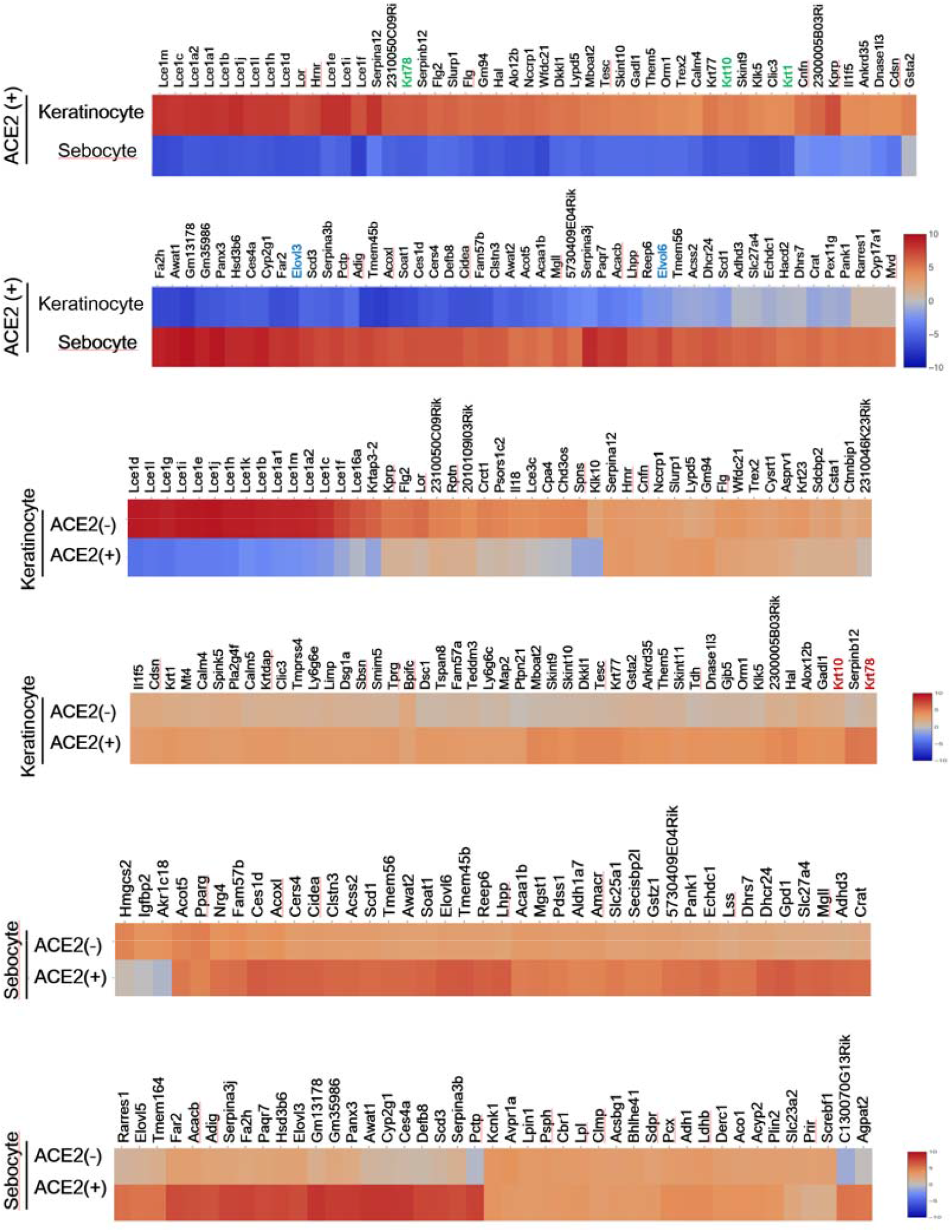
Highly different gene expression in both keratinocyte and sebocyte, further analysis the ACE2 positive and negative subset, spatial specificity genes, including krt10, krt78, krt1 etc, further implication of ACE2 expression related to location in full-thickness and degree of maturation in murine skin healing process.

**Figure 3.**
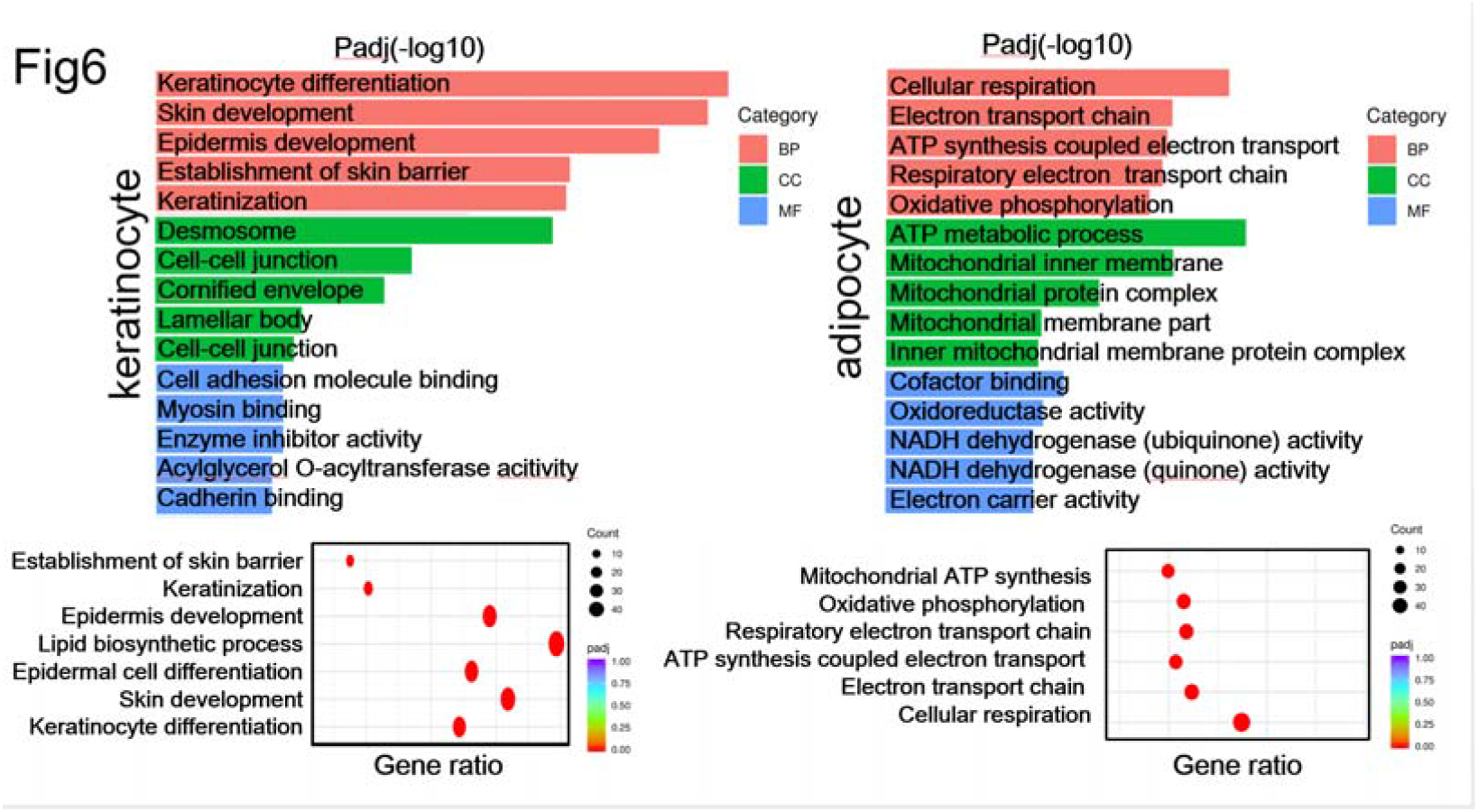
KEGG and GO analysis implication of the function and signaling pathway in different cell types highly expressed ACE2.

**Figure 4.**
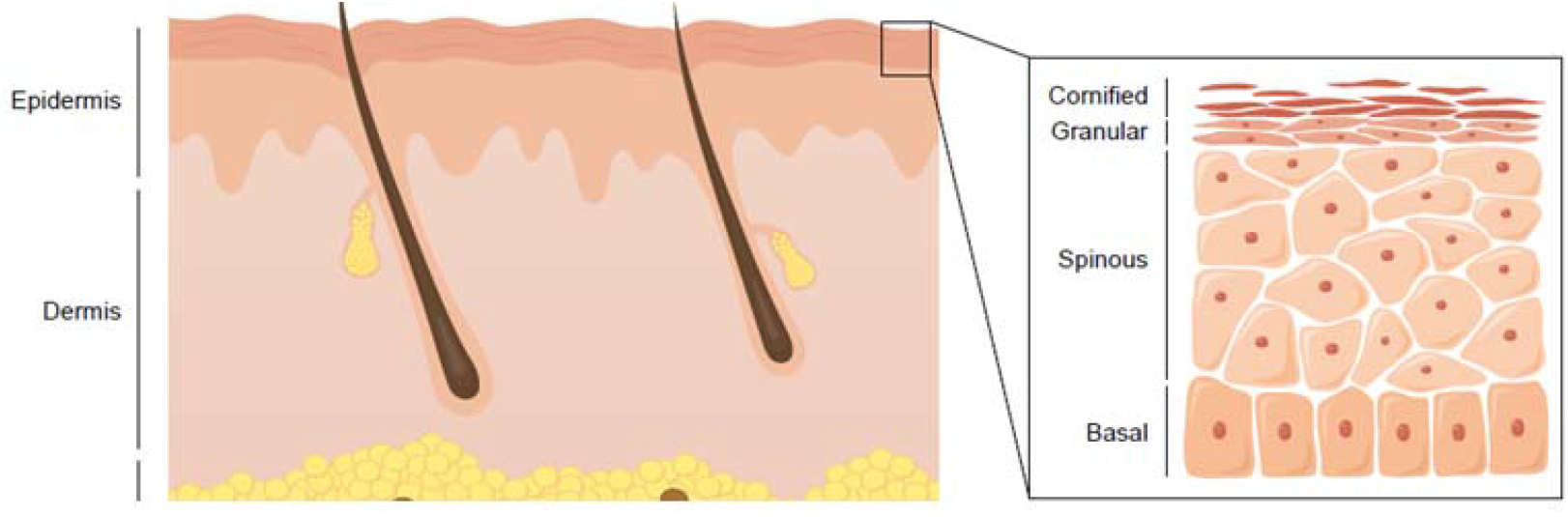
Spatial specific gene expression in different layer of murine skin, krt1 and krt5 which presented the early stage of keratinocyte maturation, i.e., which is low-differentiation keratinocyte, highly expressed in basal layer. However, the classic late-stage differentiation marker of keratinocyte, krt10 or newly reported specific marker krt78, expressed in outer layer, including granular and comified layer.

**Figure 5.**
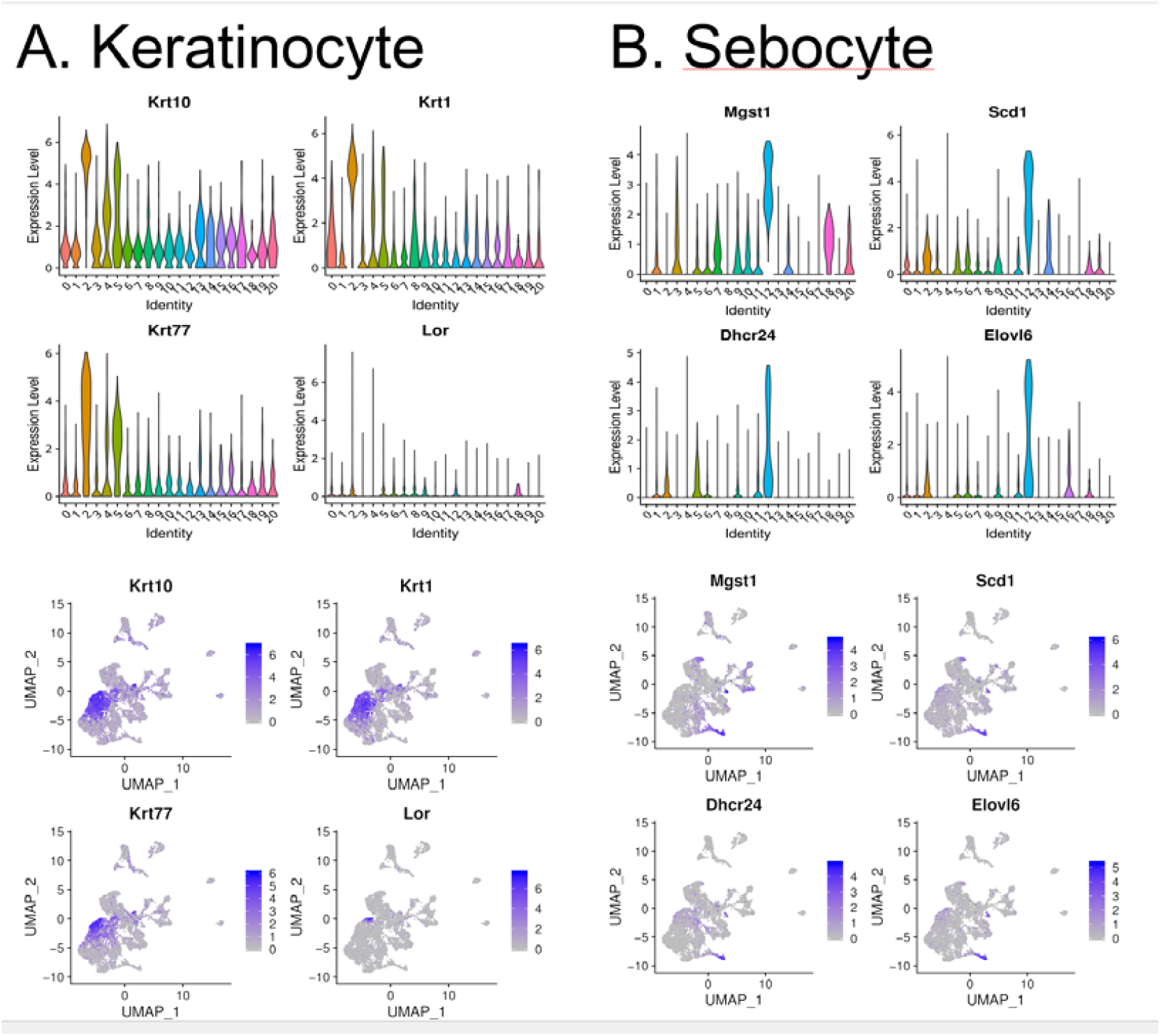
Top 4 high expression gene in different cell type, and the expression area in UMAP.

## 4. Discussion

Gene enrichment in these cellular metabolic processes indicates active energy-consuming processes in sebaceous gland after skin wounding. This might be related to function of sebocytes. Mouse sebocytes produce sebum (containing triglycerides, diglycerides, and free fatty acids, wax esters, and cholesterol) in a holocrine approach, which is crucial for hair growth, moisturization of skin and hair, and the prevention of water evaporation from the skin surface [14, 15]. The sebaceous gland arises as an outgrowth of the outer root sheath of the hair follicle as undifferentiated sebocytes emerge from peripheral cells and then move centrally (central cells) as first partially, and later fully, differentiated sebocytes in fig.4 [16]. The cuboidal or flattened peripheral cells contain no lipids, while the matured central cells have a frothy appearance as a result of the accumulation of cytoplasmic lipids.[14] In cluster 12 (sebocyte), Ace2+ cells (26%) showed up-regulated genes including stearoyl-CoA desaturase 1 (Scd1), Scd3, cell death-inducing DNA fragmentation factor (DFFA)-like effector (CIDE) a (Cidea), elongation of very-long-chain fatty acid protein 3 (Elovl3), and phosphatidylcholine transfer protein (Pctp), in comparison with Ace2^+^ cells in fig.6 (74%). Among them, Scd1 and Scd3 are specifically expressed in the sebaceous gland of mouse skin [17, 18]. Previous studies demonstrated that Scd1 expressing sebocytes derive from precursor cells mobilised from the hair follicle bulge. The precursor cells are marked by expression of Lrig1 [19]. When we overlap cells that respectively express Lrig1, Scd1 and Scd3 with Ace2^+/-^ cells in Umap, we found that the Lrig1 is more sufficiently expressed in Ace2^-^ cells (Fig. ?), suggesting that these cells might be located at peripheral zones in the sebaceous gland. Among the up-regulated genes in Ace2^+^ cells, protein of Elovl3 and Scd1 regulate lipid synthesis. Cidea is a crucial regulator of sebaceous gland lipid storage and sebum lipid secretion [20]. These functional genes are up-regulated in sebocytes expressing Ace2, which further indicate that they are matured central cells in sebaceous gland. We also observed that sebocytes resemble adipocytes in gene expressions despite their differences in lipid production, composition, and release. The afore-mentioned Elovl3 and Cidea were also marker genes for brown adipocytes. Pctp is up-regulated in Ace2^+^ cells, and has a critical role in regulating mitochondrial oxidation of fatty acids in metabolically active tissues, e.g. it is crucial for regulating adaptive thermogenesis in brown adipose tissue (BAT) [21-23]. Ace2^-^ cells were featured by increased expression of mitochondrial 3-hydroxy-3-methylglutaryl CoA synthase 2 (Hmgcs2) and insulin-like growth factor (IGF) binding-protein 2 (Igfbp2). Hmgcs2 has been implicated in the synthesis of ketone bodies, which can regulate adipocyte browning [24]. It also contributes to cellular hydroxymethylglutaryl-coenzyme A (HMG-CoA) levels and thereby provides substrate for mevalonate synthesis, the inhibition of which affects uncoupling in mature human and murine brown adipocytes, mitochondrial respiration, and browning of the white adipocytes [25].The Igfbp2 protein is the predominant Igfbp produced by white preadipocytes during adipogenesis [26]. The level of Igfbp2 expressed by adipocytes was associated with fat mass, while circulating Igfbp2 was positively associated with insulin sensitivity [27, 28]. When assessing the effect of an epidemic, transmissibility and severity are the 2 most critical factors [29]. An estimated basic reproductive number (R0, a commonly used measure of transmissibility and is defined as the number of additional persons one case infects over the course of their illness) is 2.2, which indicates the infection has the potential for sustained transmission [30]. However, the transmission routes are still not fully clarified. As a target for SARS-CoV-2, the distribution of Ace2 might offer clues for exploring possible transmission routes. In the mouse wounded skin sample, cells that express Ace2 are suprabasal keratinocytes in the epidermis and sebum-secreting sebocytes in sebaceous glands. Whether the pathogen can infect the organism through skin contact, especially when the skin is wounded, remains to be verified.

## Funding information

This work was supported by Research and Develop Program, West China Hospital of Stomatology Sichuan University (No.LCYJ2019-19); the Fundamental Research Funds for the Central Universities (No. 2082604401239)

## Statement of conflict of interest

There are no conflicts of interest related to this manuscript.

## Figure legends

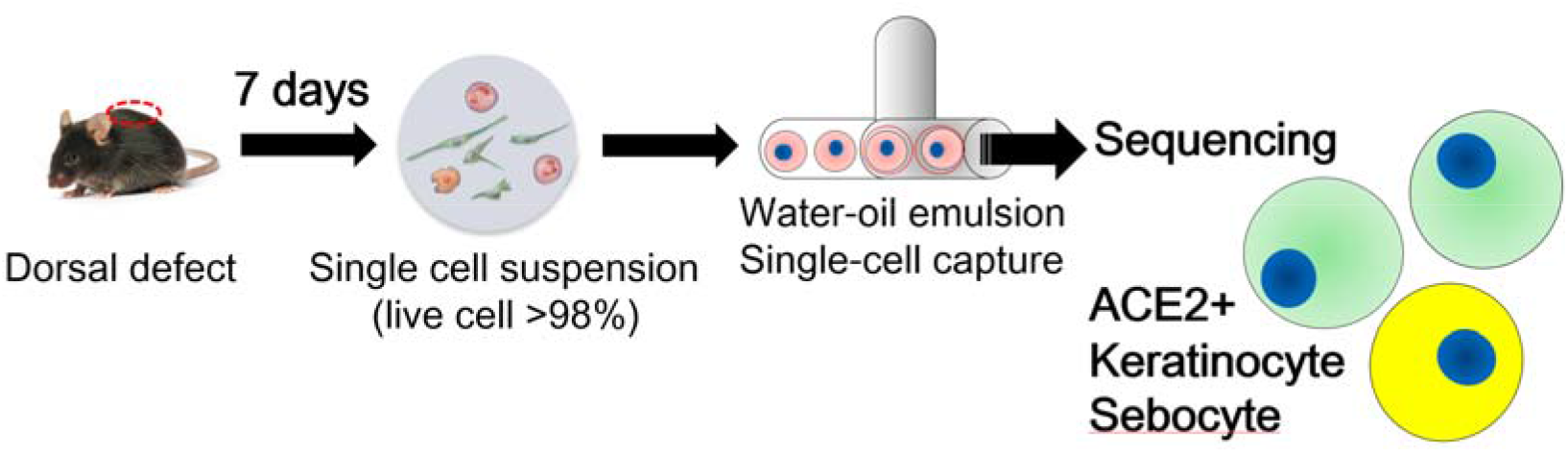

**Illustration of whole experimental design and cell-clustering process under single cell RNA-seq level**

**Illustration of experiments**

## Notes

### Competing Interest Statement

The authors have declared no competing interest.

